# The effects of *in-vitro* pH decrease on the gametogenesis of the red tree coral, *Primnoa pacifica*

**DOI:** 10.1101/407403

**Authors:** Ashley M Rossin, Rhian G Waller, Robert P Stone

## Abstract

*Primnoa pacifica* is the most ecologically important coral species in the North Pacific Ocean where it provides important habitat for commercially important fish and invertebrates. Ocean acidification (OA) is more rapidly increasing in high-latitude seas because anthropogenic CO_2_ uptake is greater in these regions. This is due to the solubility of CO_2_ in cold water and the reduced buffering capacity due to low alkalinity of colder waters. *Primnoa pacifica* colonies were cultured for six to nine months in either pH 7.55 (predicted 2100 pH levels) or pH 7.75 (control). Oocyte development and fecundity in females, and spermatocyst stages in males were measured to assess the effects of pH on gametogenesis. Oocyte diameters were 13.6% smaller and fecundities were 30.9% lower in the Year 2100 samples, indicating that OA may limit oocyte formation, potentially through lipid limitation. A higher proportion of vitellogenic oocytes (65%) were also reabsorbed (oosorption) in the Year 2100 treatment. Lowered pH appeared to advance the process of spermatogenesis with a higher percentage of later stage sperm compared to control controls. There was a laboratory effect observed in all measurement types, however these only significantly affected the analyses of spermatogenesis. These results indicate that reproduction may not be possible in an acidified ocean, or that if spawning could occur, spawned oocytes would not be sufficiently equipped to support the normal development of larvae.

## Introduction

Primnoidae is one of the most abundant gorgonian families in deep-sea and polar regions [1] and the most abundant coral family in Alaskan waters [2]. *Primnoa pacifica* Kinoshita, 1907 or red tree coral is one of the most common habitat forming and ecologically important coldwater octocorals in the Northeast Pacific Ocean [3-6]. It has a broad geographic range from the Sea of Japan and Sea of Okhotsk, through the Aleutian Islands and Gulf of Alaska (GOA), and to British Columbia and is found in depths from 6–573 m [7]. The shallowest depths recorded are from the GOA where the phenomenon of deep-water emergence occurs in the glacial fjords [8]. These corals prefer habitats dominated by sloping bedrock on rough seabeds and areas with moderate water currents [4].

*Primnoa pacifica* exhibits keystone species characteristics [4] as described by King and Beazley [12] and are undoubtedly foundation species [Powers 1996] in the habitats were they form dense thickets in the Gulf of Alaska. The dense thickets in the GOA provide essential habitat for economically important species [4,8,10,11]. *P. pacifica* is also an ecosystem engineer, modifying its environment by altering small-scale ocean currents and creating living spaces for other organisms [8,9], highlighting this species’ importance to the GoA and shelf fjord ecosystems.

Anthropogenic activities, including ocean acidification (OA), may affect primnoids and other calcifying organisms. As a result of increased atmospheric CO_2_, average surface ocean pH has decreased by 0.1 pH units since the Industrial Revolution and is projected to decrease by another 0.3–0.4 units by the end of the century [13-16]. The saturation horizons of calcite and aragonite (CSH and ASH, respectively) in the North Pacific Ocean are naturally shallow (~200 m) relative to other oceans [Jessica Cross, NOAA Pacific Marine Environmental Laboratory, pers. comm.] and are shoaling at a rate of 1–2 m per year [17]. Recent studies have suggested that calcifying organisms at high latitudes are at immediate risk from OA due to seawater being only slightly supersaturated with regard to calcium carbonate [18,19].

Increased pCO_2_ may have complex effects on the physiology, growth and reproductive success of marine calcifiers [20]. A greater portion of an organism’s energy budget may be partitioned towards maintenance of the acid-base status of internal fluids and away from other fitness-sustaining processes such as shell and somatic growth, immune response, protein synthesis, behavior, and reproduction [21]. OA has the potential to negatively impact sexual reproduction through development of multiple early life stages and may contribute to substantial declines in recruitment that will cascade from community to ecosystem scales [15,22].

Red tree corals exhibit carbonate polymorphism, ranging from 65.6% aragonite (axes) to 99.7% high magnesium calcite (sclerites), the most soluble form of calcium carbonate [R. Stone unpublished data]. Some organisms may be able to upregulate pH and continue to grow or calcify under acidic conditions, however, this might come at the cost of affecting other physiological processes, such as reproduction [23]. While corals might physically appear healthy, metabolic costs limiting reproduction could be deleterious to the population. The objective of this study was to experimentally investigate the effects of OA on gamete production of *P*. *pacifica*.

## Materials and methods

Ethical approval for this research was not required by any federal, state, or international law because the animals used were invertebrates. The transportation and field collection of the animals was authorized by the Alaska Department of Fish & Game (Fish Resource Permit CF16-027). Reference to trade names does not imply endorsement by the National Marine Fisheries Service, NOAA.

### Sample Collection Area

Samples for this study were collected from a single site in Tracy Arm fjord, Holkham Bay in Southeast Alaska (Fig 1). The glacial fjord is 49 km long, up to 378 m deep and terminates at two tide-water glaciers, the Sawyer and the South Sawyer. Previous surveys of the fjord revealed thickets of *P*. *pacifica* as shallow as 6 m and to depths greater than 100 m [R. Stone, unpublished data]. The collection site was located in the central part of the fjord, 13 and 15 km from the two tide-water glaciers respectively.

**Figure 1.**
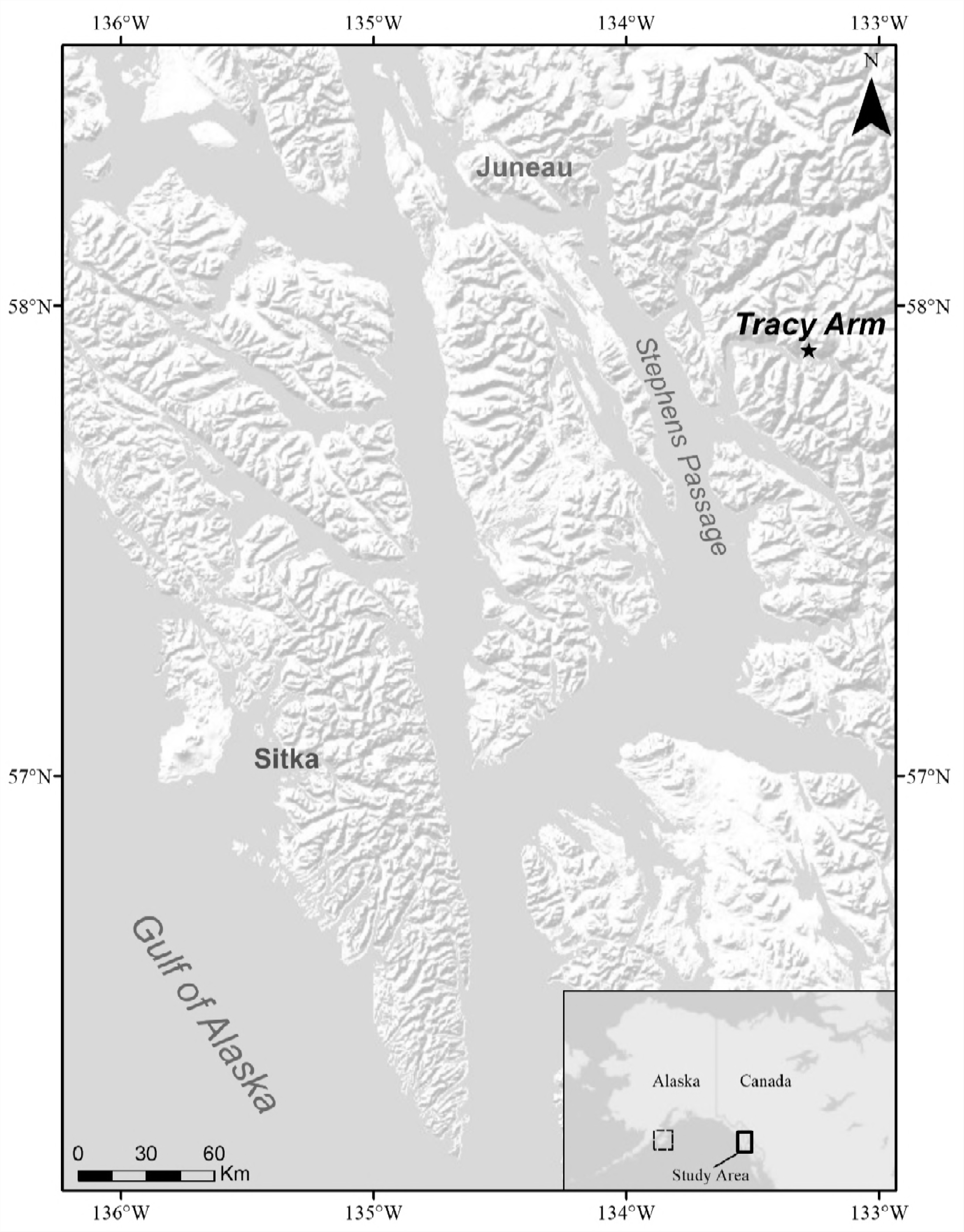
Sample collection and experimental laboratory site. The sample collection site in Tracy Arm is denoted by the star. In the inset map, the solid box indicates the zoomed in region and the dotted box marks Kodiak Island, where the laboratory experiment was completed.

From January 8–11, 2016, 54 *P*. *pacifica* colonies with healthy, intact growing tips were sampled with SCUBA at depths between 10–19 m. Colony height was measured and three sprigs, 10–15 cm long, were sampled from each colony (Fig 2). One sprig from each colony was immediately fixed in a 4% borax buffered formalin solution for 24 hours, transferred to 70% ethanol, and shipped to the Darling Marine Center (Walpole, Maine, USA) for histological processing. The other two sprigs were maintained live in circulating ambient seawater until they were transported in 250-ml Nalgene containers of ambient seawater, saturated with oxygen, packed on blue ice, and transported in coolers via commercial airliner to the NOAA Kodiak Laboratory (Kodiak, Alaska, USA; Fig 1). Sex was already known for more than half of the experimental colonies [8]; sex for the remaining colonies was determined during collection and later reconfirmed by histology in the laboratory. The colonies sampled consisted of 30 females, 20 males, and 4 were non-reproductive.

**Figure 2.**
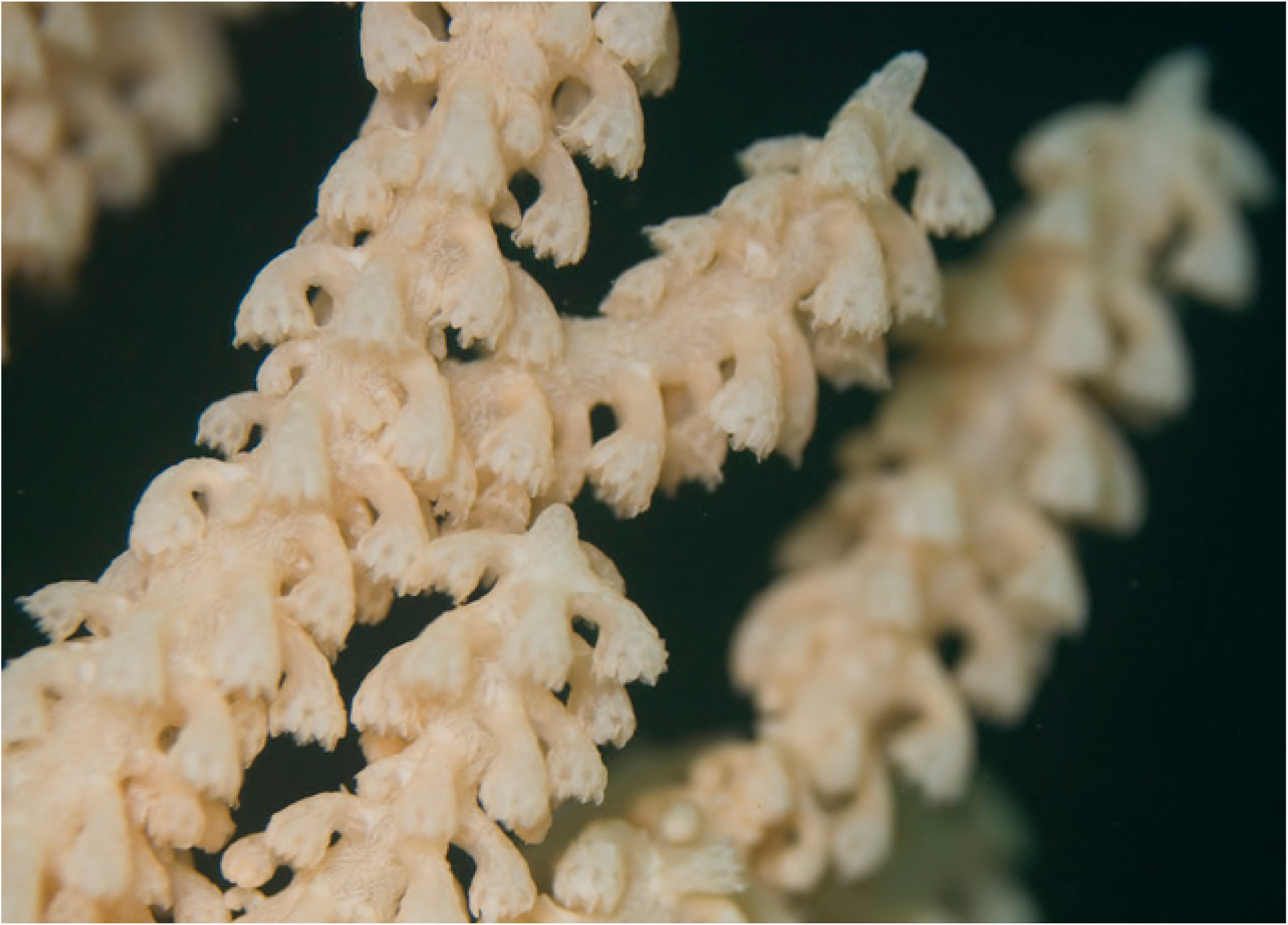
Image of growing tip on a *Primnoa pacifica* colony *in situ*.

### Laboratory Experiment

The design for this experiment consisted of three replicates of two pH treatments: One sprig from each colony was randomly assigned to two experimental treatments: (1) Year 2100, which is the predicted pH at a depth of 175 m in the eastern GOA in 2100 (7.55 pH units) and (2) Control, which was the pH (7.75 pH units) for the same region in 2016. The projected Year 2100 pH following RCP8.5 (high scenario) is from the Intergovernmental Panel on Climate Change (IPPC) as used in the Community Earth System Model 4 [14; CESM4; Jessica Cross, NOAA Pacific Marine Environmental Laboratory, pers. comm.]. Treatment aquaria measured 120 cm (L) X 60 cm (W) X 60 cm (H) and unfiltered seawater was pumped into the laboratory from Trident Basin (~20.5 m depth) to a head tank and then delivered at 2 l/min to each experimental tank. pH was controlled with a monitored dosing system by bubbling CO_2_ directly into the experimental tanks. CO_2_ input was controlled by Honeywell controllers and Durafet III pH probes controlling a gas valve. Daily pH and temperature measurements were made in each tank. Once a week, water samples were collected from each tank, fixed with 0.02% mercuric chloride, and sent to the University of Alaska Ocean Acidification Research Center for alkalinity and dissolved inorganic carbon (DIC) analysis using standardized methods [25]. Those measured results were used to calculate pH, pCO2, HCO3^-^, CO3^-2^, Ω_aragonite_, and Ω_calcite_ using the seacarb package in R [26] (R 2.14.0, Vienna, Austria).

Two sprigs from each colony were randomly assigned to a treatment and then randomly assigned to a replicate within the treatment (i.e., an aquarium) in a repeated measures design. Sprigs were kept in total darkness. Water temperature was maintained at 4.5–5°C, which is the mean annual temperature experienced by the corals *in situ* [8]. Sprigs were suspended in the water column, tip facing downwards and tied with a Spectrafiber microfilament braided line (10pound test). They were fed 25 ml of a mixture of six marine microalgae (Reed Mariculture Inc., Shellfish Diet 1800) that was diluted in 450 ml of unfiltered seawater once a week. The tanks were allowed to go static (i. e. no water flow) for 20 minutes during feeding to allow the food mixture to fully permeate the tank.

The laboratory experiment was conducted between 15 January and 22 September 2016. However, on 21 June (Day 158), the circulating water system of Tank 3 (Year 2100 treatment) failed, causing all but three sprigs to begin sloughing their tissue. Polyps from those sprigs were sampled immediately. The three remaining sprigs were sampled on 23 June (Day 160) and the corresponding sprigs for those colonies in the Control treatment were sampled on 29 June (Day 166) and immediately processed for histological analyses. The experiment was terminated on 22 September 2016 (Day 251) and tissues were prepared for histological processing.

### Histological Processing and Examination

All sprigs were assigned random numbers prior to histological processing to prevent bias. Three to nine polyps were dissected from each sprig for histological processing following previous protocols in [8]. Polyps were sampled randomly from the sprigs as previous observations have suggested no variance in gametogenesis with location in colony [R. Waller, unpublished data]. Polyps were decalcified with Rapid Bone Decalcifier (Electron Microscopy Sciences), then dehydrated in serial ethanol dilutions from 30% to 100%. Samples were then cleared in Toluene solution and then immersed in paraffin wax (Leica ParaPlast Plus) for approximately 48 hours at 56°C.

Tissue was then embedded in paraffin wax blocks and left to cool for at least 24 hours, then placed in a freezer at least one hour prior to sectioning with the microtome (Microm HM 325). All wax blocks were serially sectioned. Specimens were sliced 6 μm thick to maintain tissue quality; the distance between serial sections was 90 μm between slides, which is the average diameter of the oocyte nucleus in *P. pacifica* [8]. Sectioned tissue was mounted on glass slides, dried on slide warmers, and stained with Hematoxylin and Eosin or Masson’s Trichrome. (Fig 3).

**Figure 3.**
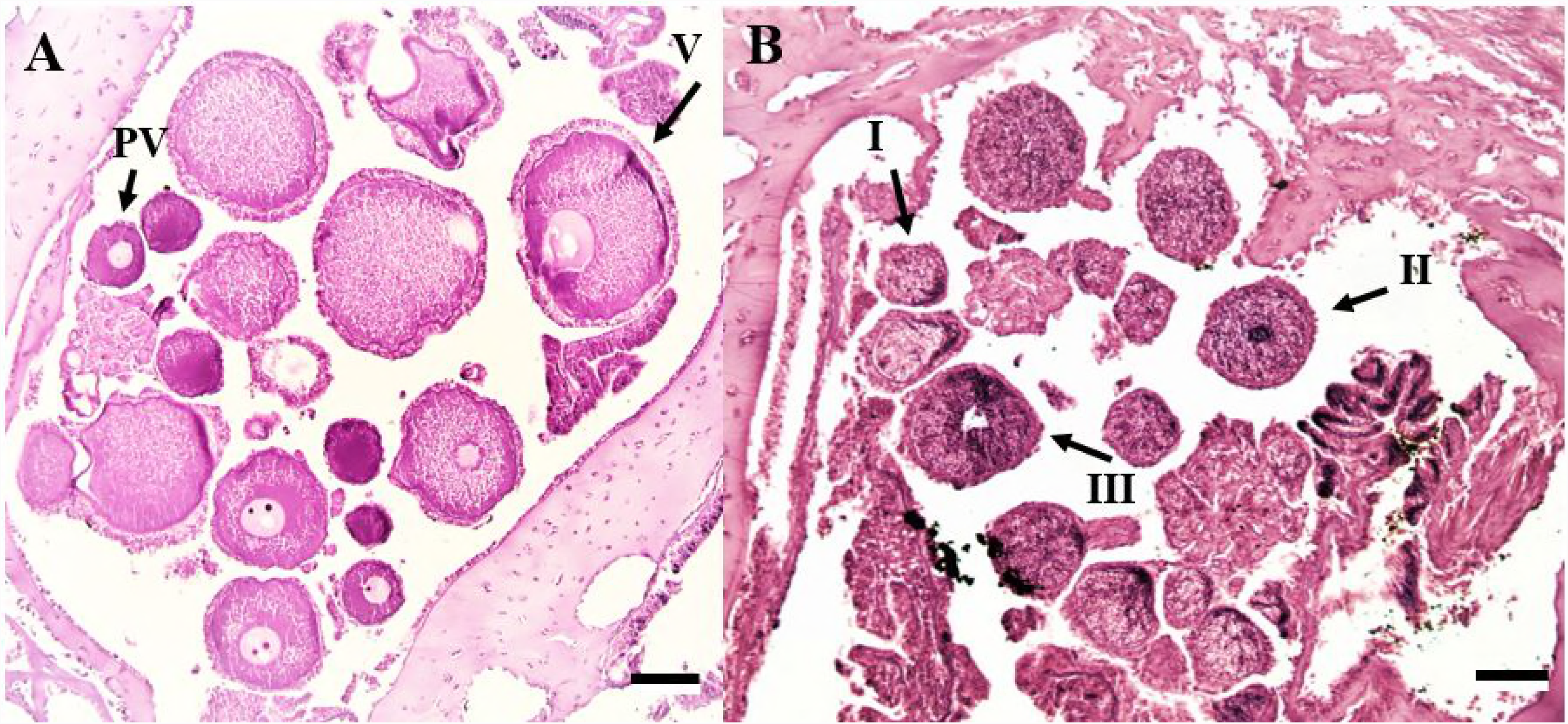
Histological sections of polyp tissue. (A) female with previtellogenic (PV) and vitellogenic (V) oocytes. (B) male with spermatocysts from three of the four stages. Scale bars represent 100 μm.

Slides were examined using an Olympus (CX31) compound microscope with a Motic video camera attachment. Images were captured using Motic Image Plus and analyzed with ImageJ (NIH) software to calculate oocyte and nucleus diameter. One hundred gametes were measured for each individual sprig.

Spermatocysts were staged from I-IV, indicating increasing maturity, following the classification by Waller et al [8]. Fecundity was measured by counting all oocytes (both previtellogenic and vitellogenic (Fig 3A) in three polyps per sprig and then averaged following Waller et al [8] to determine the average fecundity per polyp in a colony. Oocytes with a visible nucleus were the only oocytes counted to ensure there was no double counting, as the nucleus is in the center of an oocyte.

Histological analyses were initially performed using Microsoft Excel, and comparison between treatments and individuals were performed using R Studio Version 3.4.1. A Repeated Measures ANOVA (RMANOVA) was performed for all data comparisons to satisfy the “within subjects” assumption from the “car” package [26]. A paired t-test was performed within the same package to determine if the difference in variance between treatments was significant (p<0.05). The methods and results for the corresponding sprig analysis can be found in Supplementary File 1, and their corresponding p-values and results from the RMANOVA in Supplementary Appendices 1, 2, and 3.

## Results

Comparisons between this dataset and the previous work from the same coral population [8] are in the Supplementary Information (SI1) and all RMANOVA results and corresponding pvalues are in the supplementary appendix (SA1).

### Spermatogenesis

Tissue from 50 male sprigs were analyzed in this study. There were 20 sprigs from Time 0 (no experimental treatment), 18 sprigs from the Control treatment, and 12 sprigs from the Year 2100 treatment (Table 1). Males from the Time 0 subsample had a mixed composition of all four spermatocyst stages within an individual sprig. Stage II was the mode stage (Fig 4).

**Table 1.** **Number of male and female sprigs per treatment from the tank experiment.**

| *Sex* | *Time 0* | *Ambient* | *2100* |
| --- | --- | --- | --- |
| *Male* | 20 | 18 | 12 |
| *Female* | 30 | 30 | 20 |

**Figure 4.**
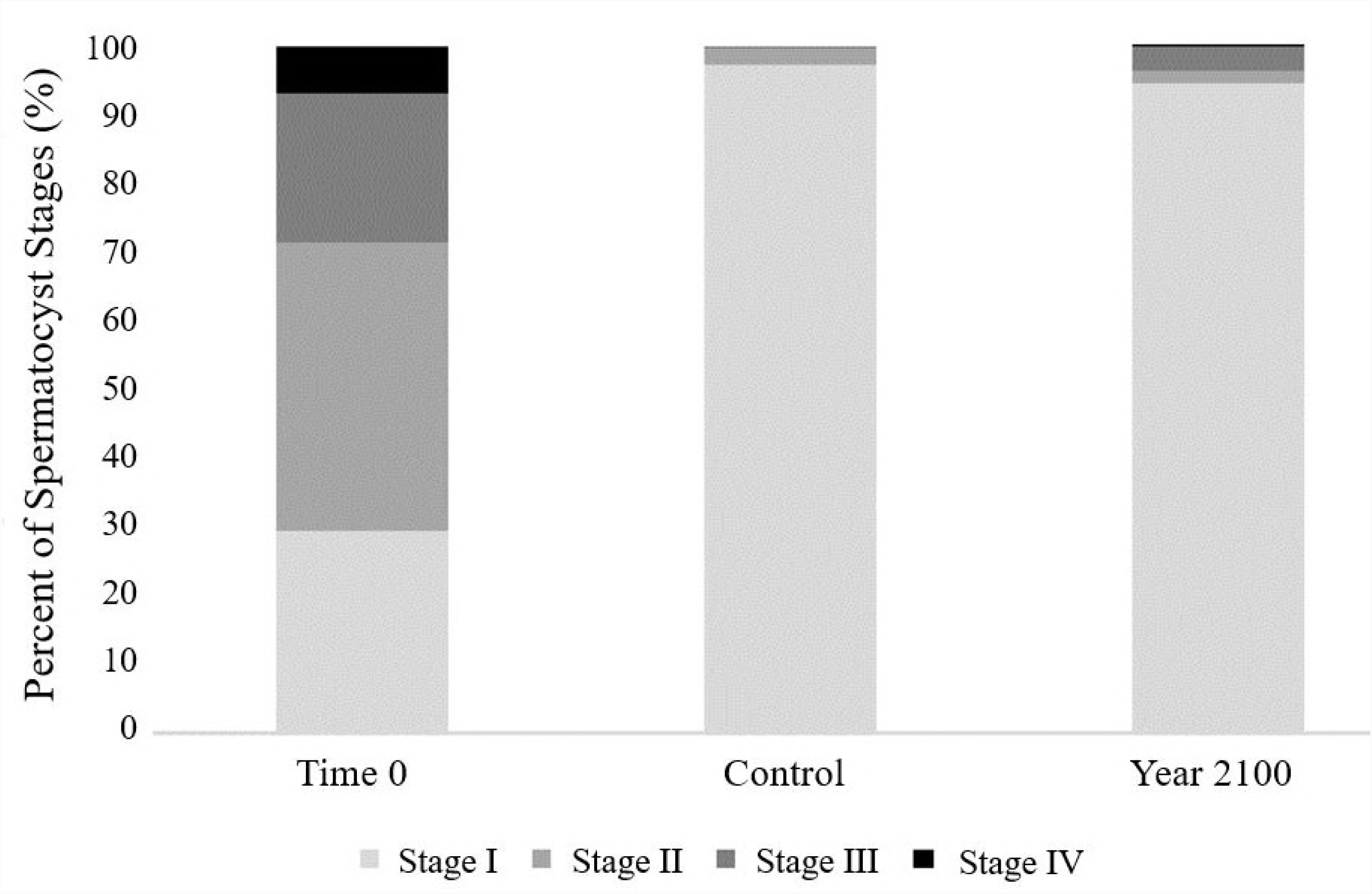
Mean percent spermatocyst stage (%) versus treatment. Sperm stages increase with maturity. N indicates number of individuals and n indicates number of spermatocysts (Time 0: N = 20, n = 2000; Control: N = 18, n = 1710; Year 2100: N = 12, n = 1070).

Males from the Control treatment show a more homogenous composition individual sprigs with 83–99% of the spermatocysts at Stage I and the remaining were Stage II or III (Fig 4). Males from the Year 2100 treatment had a slightly more varied composition with 66–99% Stage I, and the remaining were Stages II or III (Fig 4).

An RMANOVA was used to determine if there was a difference in stage composition of spermatocysts among treatments (Time 0, Control, and Year 2100). Regardless of the spermatocyst stage, treatment had an effect on mean percent of each stage (p < 0.05; SA1_1-SA1_4 Tables). Time 0 was statistically different from both Control and Year 2100 (p < 0.05; SA1_1-SA1_4 Tables), but Control and Year 2100 were not statistically different from one another (p > 0.05; SA1_1-SA1_4 Tables).

### Oogenesis

Tissue from 80 female sprigs were examined in this study (Table 1). Standard error is used as deviation from the mean. The mean oocyte diameter for Time 0 was 106.8 ± 1.09 μm, Control was 77.79 ± 1.12 μm and Year 2100 was 67.18 ± 1.09 μm (Fig 5). Mean oocyte diameters significantly differed among the three treatments (RMANOVA, p < 0.05; SA1_5 Table) and each treatment was statistically different from one another (RMANOVA, p < 0.05; SA1_5 Table).

**Figure 5.**
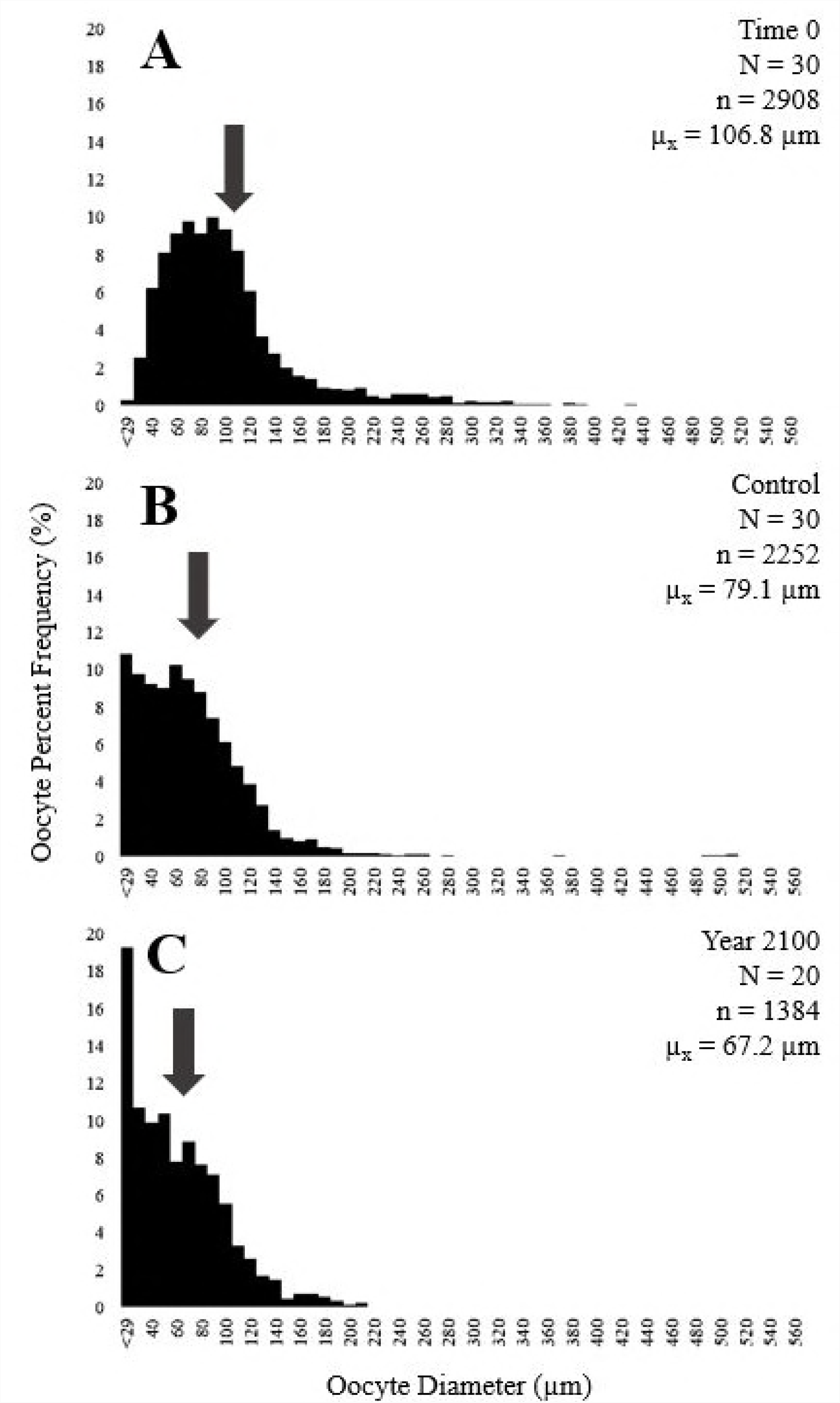
Histogram of percent oocyte frequency (%) versus oocyte diameter (μm). (A) Time 0, (B) Control, and (C) Year 2100. N indicates number of individuals, n indicates number of oocytes measured, and μ_x_ is the mean oocyte diameter (indicated by the arrows).

### Oosorption

Oosorption is the process of resorbing vitellogenic oocytes to use the lipids for other processes when under stress [27]. Thirty eight of the 80 female sprigs (48%) had structures near oocytes which did not have a nucleus and were composed of lipid-dense concentrations (Fig 6). These would have been the approximate size (~220–802 μm) of vitellogenic oocytes [8] if a nucleus had been present and were observed alongside both previtellogenic and vitellogenic oocytes. Twenty percent of the Time 0 female sprigs, 63% of the Control female sprigs, and 65% of the Year 2100 female sprigs had these structures. There was no apparent relationship between the presence of cells undergoing oosorption and treatment type nor was the presence of the structures at Time 0 an apparent indicator for presence in Control or Year 2100 sprigs from the same colony.

**Figure 6.**
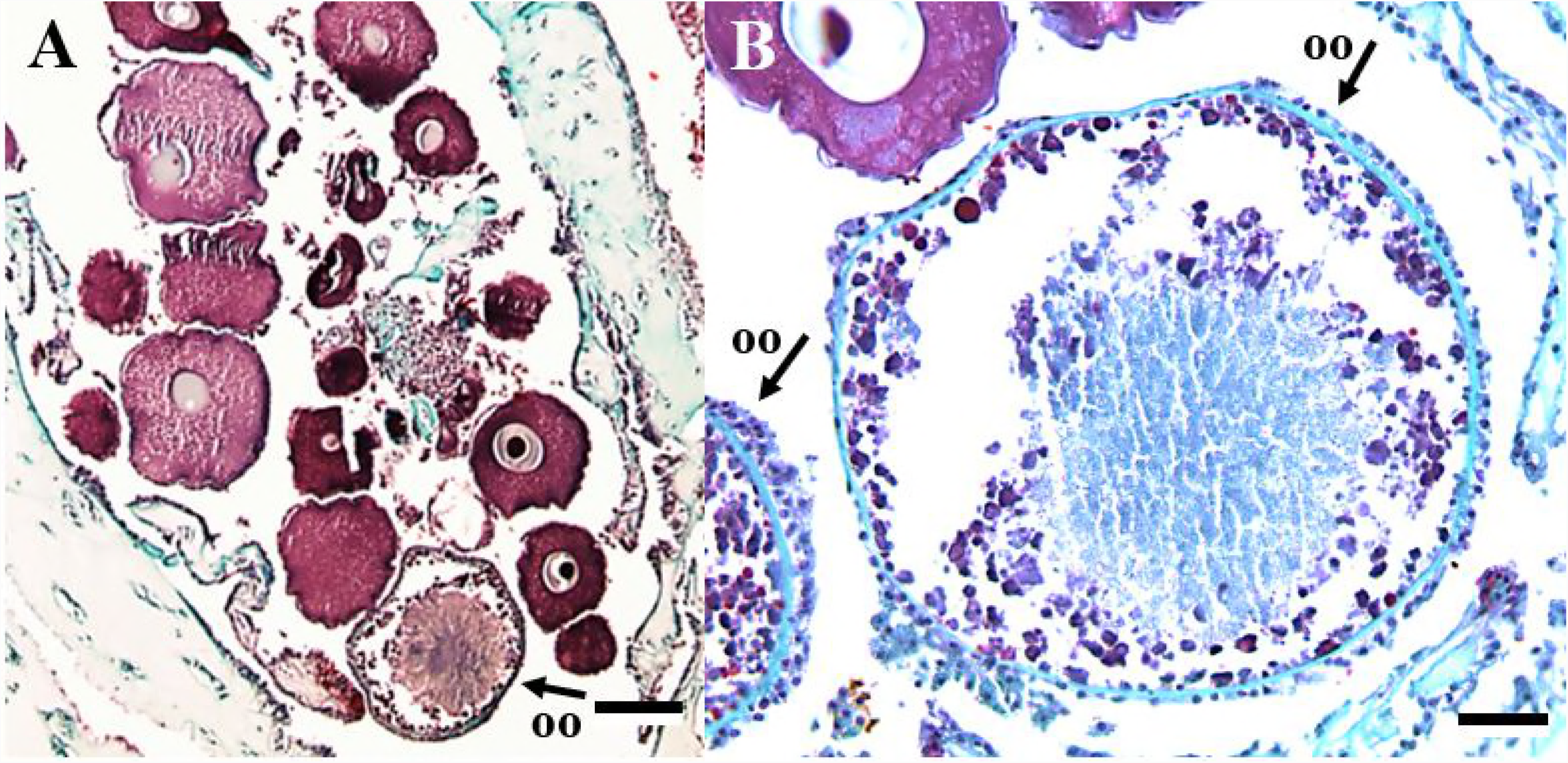
Light microscopy photograph of female polyp containing a non-nucleated oocyte. (A) shows oosorption in polyp at 4X magnification. (B) shows oosorption at 10X magnification. Scale bars are 100 μm and 500 μm respectively. OO = oocyte undergoing oosorption.

### Fecundity

All reproductive females had a maximum colony height between 42 and 160 cm and maximum fecundity was positively related to colony height, albeit with a low coefficient of determination (R^2^=0.06; Fig 7).

**Figure 7.**
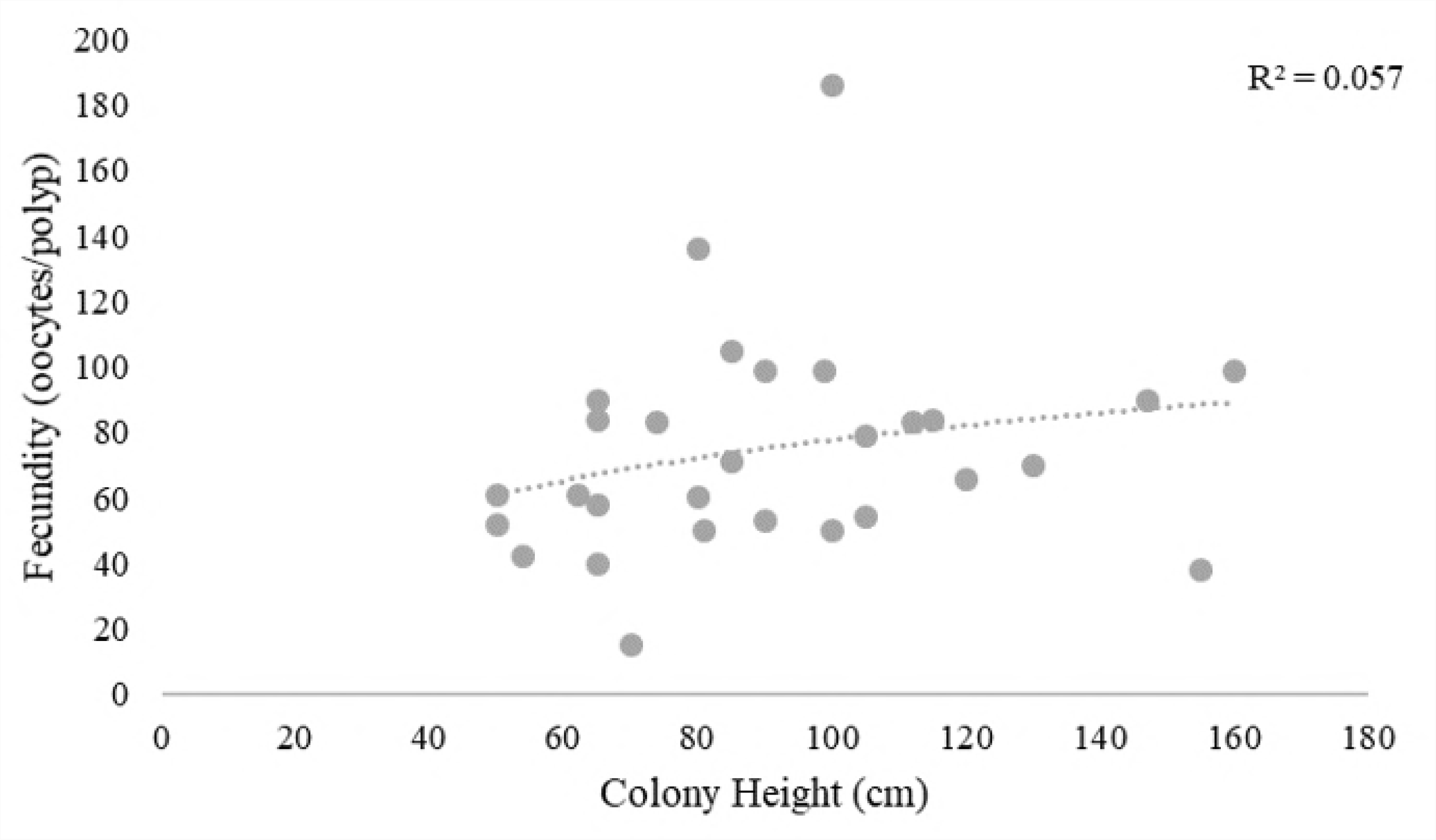
Maximum fecundity (oocytes per polyp) versus colony height. Trend line is logarithmic.

The average (SE) fecundity for Time 0 was 56.43 ± 3.13 oocytes per polyp, Control was 25.1 ± 3.19 oocytes per polyp, and Year 2100 was 17.32 ± 2.24 oocytes per polyp (Fig 8). Treatment had an effect on average fecundity (RMANOVA, p < 0.05; SA1_6 Table), and each treatment was statistically different from the others (RMANOVA, p < 0.05; SA1_6 Table).

**Figure 8.**
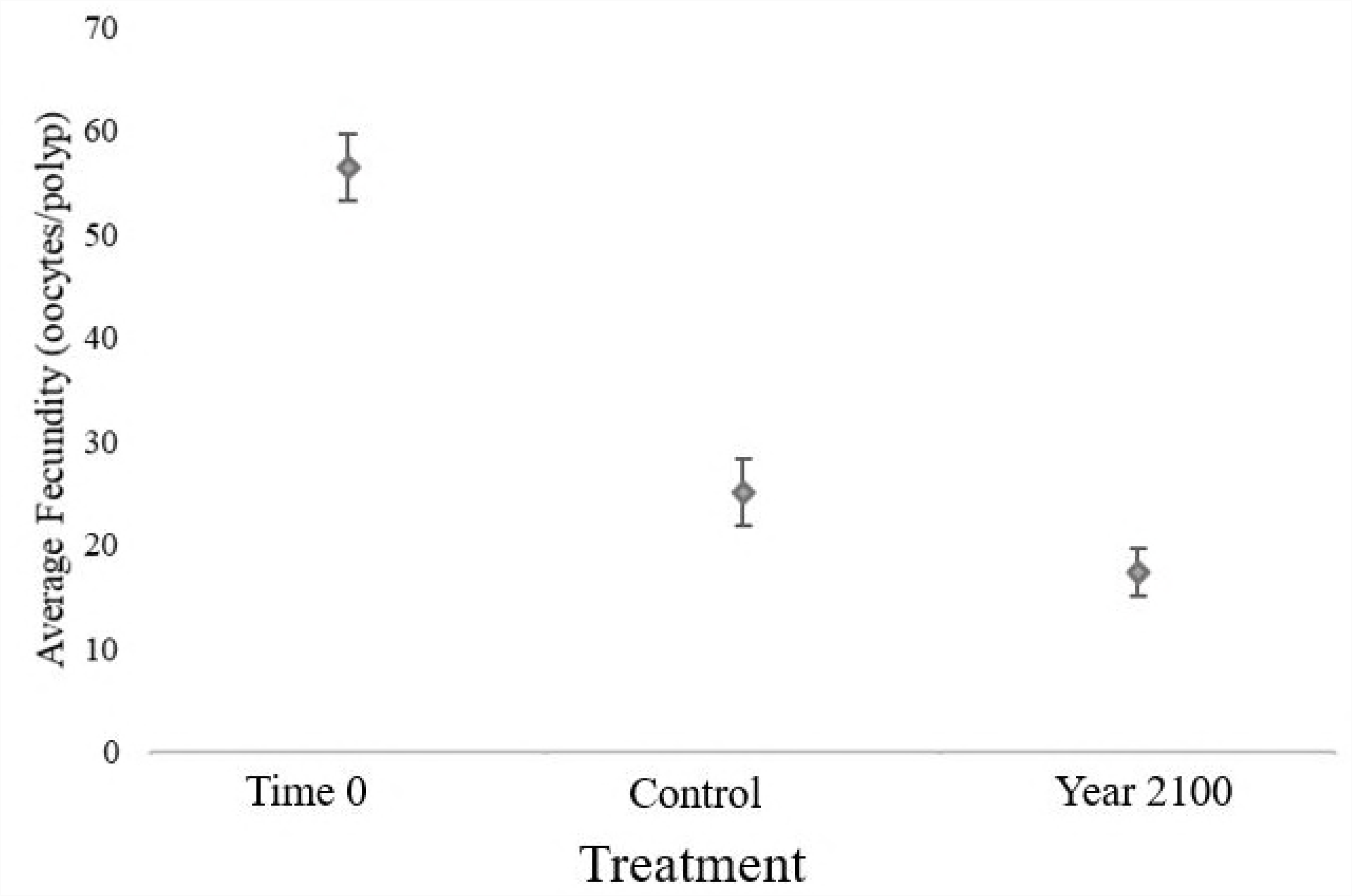
Average fecundity (oocytes per polyp) for sprigs versus treatment. Error bars indicate standard error. N represents number of individuals (Time 0: N = 30; Control: N = 30; Year 2100: N = 20).

## Discussion

The sprigs from the Control and Year 2100 treatments were significantly different than the Time 0 samples. The Control and Year 2100 results were statistically the same for the spermatocyst stages, while the oocyte diameters and fecundities were statistically different among all treatments. This negative trend in fecundity and development could be attributed to the animals being stressed by being in the tanks, regardless of treatment, and likely reflects the absence of a homogeneity between the tank environment and the corals’ natural habitat [28].

Though the laboratory effect did cause significant differences in oocyte diameters and fecundities; effects from the pH treatment can still be discerned. Females in the Year 2100 treatment had a smaller average oocyte size and lower fecundity than those in the Control treatment. The Year 2100 females also had the highest proportion of resorbing oocytes compared to Control and Time 0. Considering these differences, it appears improbable that these females could successfully reproduce in the projected pH for Year 2100. Oocytes would likely be too small to fertilize successfully, and even if some did produce larvae, fecundities might be too low for effective recruitment events.

### Spermatogenesis

The Time 0 samples were added to the existing reproductive dataset from Waller et al [8], and although the exact composition shifts each year and month, in all January time points, all four spermatocyst stages were present (Supplementary Information 1). The Time 0 samples had a heterogeneous composition, the Year 2100 treatment had a slightly varied composition, and the Control treatment was composed almost entirely of Stage I.

While there was no significant negative difference between Control and Year 2100 treatments with regards to spermatocyst composition, the Year 2100 spermatocysts were more developed than those from Control. There is still much to learn regarding spermatogenesis and OA as well as the additive effects of acidified conditions on fertilization. Any improved potential in sperm condition could potentially be cancelled out by reduced oocyte fitness, fertilization success, or embryonic condition [29,30].

### Oogenesis

The average oocyte diameter for Time 0 was 106.8 μm, Control was 77.8 μm, and Year 2100 was 67.2 μm (Fig 5), which were all significantly distinct from one another (p < 2.2 e-16). While differences in oocyte diameter between Time 0 and both treatments indicate the shift in seasonality and laboratory effect, the difference between the Control and 2100 may indicate a more notable trend. The Year 2100 oocytes are smaller than those from Control, indicating less lipid reserves available for gametogenesis and reproduction which can impact multiple life history stages. For example, smaller eggs in sea urchins under stressed conditions can have less lipid content [31,32] with negative consequences for larval fitness such as increased age at metamorphosis and reduced fitness of newly settled juveniles [32]. Larval longevity is also dependent on high lipid content of oocytes in some reef building scleractinian corals [33-36].

### Oosorption

The combined results of smaller average oocyte diameters for Year 2100 compared to Control treatments (67.2 μm and 79.1 μm, respectively) and an increased presence of large, non-nucleated lipid-dense structures (65% and 63%) suggest that lipid reserves may have been redirected from gametogenesis to other metabolic processes under lower pH conditions and lab conditions. The increased occurrence of oosorption coinciding with smaller oocytes could indicate that spawning of fully developed oocytes, or natural spawning may not be possible under acidified conditions.

The lipid-dense structures were observed within reproductive polyps, alongside both previtellogenic and vitellogenic eggs, and generally near the gastrovascular wall. Initially, we considered the structures to be unidentified extracellular material, however they are more consistent with oosorbed oocytes when compared with previous observations [27]. These structures had a membrane surrounding the mass of cells, similar to the vitelline envelope in oocytes (Fig 6). Small cells surrounding the membrane appeared to have suspended nuclei which were connected by a less rigid conglomerate, similar to previous observations [27] of intracellular gaps leading to an unorganized association of cells on the exterior of the oocyte.

Small, round concentrations of cells within the membrane stained as lipids, as oocytes do, however, they were more densely packed and unorganized compared to those in vitellogenic oocytes.

In this study we observed no relationship between fecundity and oocyte size with the presence of oosorption, so oosorption does not prevent other oocytes from developing within the same individual, and smaller eggs are likely held in reserve to develop from the resorbed lipids [27]. The process likely occurs naturally since these structures were observed in females from all treatment groups; and may not be unique to *P. pacifica* or gorgonians. *Acropora millepora* uses lipid reserves when under stress to provide energy to maintain net calcification rates [37].

The increase in oosorption rate in experimental treatments compared to ambient Day 0 likely indicates a stress response of the corals to laboratory and to a lesser degree low pH conditions. Oosorption has been indicated as a stress response to adverse holding conditions including inadequate food supply or poor quality, stagnant water supply, and varying water temperatures [27]. Species without significant nutrient reserves that devote a large portion of their energy budget to gametogenesis may also resorb gametes in response to starvation or stressors that induce a rapid energy deficit [27].

### Fecundity

The fecundity and height relationship from this study corroborates previous results for *P. pacifica* that female colonies with total height greater than 50 cm are reproductive [8], and that there is a slightly positive relationship with size and fecundity as shown by the logarithmic line (Fig 7), all non-reproductive individuals were smaller than 90 cm (SP1 Table).

While all three fecundities are statistically different from one another, the Year 2100 average fecundity is significantly lower than the Control treatment. As fecundity can be used as a proxy for reproductive effort of a colony [38], this marked decrease in fecundity with an increase in acidity elucidates a decrease in reproductive effort under stress.

## Conclusions

The results from this study are just the beginning for examining the effects of OA on the allocation of energy using the processes of gametogenesis and reproduction as proxies for coldwater corals. While the spermatogenesis results cannot be fully compared due to the observed laboratory effect, the oogenesis results are notable. The smaller and more infrequent occurrence (lower fecundity) of oocytes from the Year 2100 females indicates an inability to fully allocate resources to oogenesis in acidified conditions. These results combined with the increased presence of oosorption could have potentially deleterious effects for red tree coral populations in future oceans if OA continues at projected levels.

The apparent laboratory effect observed in this study indicates a need to better understand the natural environment of cold-water corals, particularly gorgonians to more accurately replicate *in situ* conditions in laboratory experiments. Though our goal was to conduct the experiment for one year, or a full reproductive cycle, we were unable to maintain the corals for more than 200 days. We recommend that future experiments with red tree corals be limited to 200 days to avoid the deleterious physiological effects of residing in the laboratory or alternatively that in *situ* free-ocean CO_2_ enrichment experiments be used [39].

While commercially important species have been the primary focus of OA studies to date, there is a need to understand the effects of OA on other species that play other important roles in marine ecosystems. Keystone and foundation species that create habitat and provide structure for commercially important species need to be included in the portfolio of this emerging area of research and should include multiple life history stages to understand potential carry-over effects from generation to generation.

## Acknowledgements

We thank Bob Foy, Allie Bateman, Nikki Gabriel, and Chris Long (NOAA’s Kodiak Fisheries Research Center) for assistance setting up and maintaining the experiments. We also thank Captain Dan Foley and the crew of the MV *Steller* for field support and Elise Hartill for help collecting the corals. Thank you to Genny Wilson for staining the slides for this project. Thank you to Robert Steneck and Aaron Strong for your edits on this work. We also thank Michele Matsuda for your statistical help. We thank Bob Foy for reviewing an earlier draft of this manuscript. This project was funded by NOAA Fisheries (Alaska Fisheries Science Center) and NOAA’s Ocean Acidification Program. The findings and conclusions in the paper are those of the authors and do not necessarily represent the views of the National Marine Fisheries Service.

## Supporting Information

**SP1. Supplementary protocol 1.** Information detailing how the corals were arranged in the tanks and protocol for their care for the duration of the OA experiment.

**Figure SP1. Tank setup at Kodiak Fisheries Research Center, AK.** Tanks 1, 5, and 6 were treatment A and tanks 2, 3, and 4 were treatment B. Each tank held 18 sprigs split between four rows. Highlighted cells represent individuals that were “spares” as their sex could not be determined to absolute certainty on the cruise.

**Table SP1.** Colony height and sex of collected colonies.

**SF1. Supplementary file 1.** Methods and results of the sprig analysis.

**Figure SF1**. **Boxplot distribution of percent change (%) versus treatment and measurement type.** N represents number of individuals. For each treatment and methodology, N = 35.

**Table SF1. Average results from exterior sprig measurements.** For all measurement types and sampling dates, the number of sprigs was 35. SE indicates standard error.

**SI1. Supplementary information 1.** The OA study was compared to the 2014 reproductive study by Waller and associates.

**Table SI1. End of experiment average oocyte diameter split by collection month.** SE indicates standard error.

**Table SI2. Average oocyte diameters for 2014 reproductive dataset.** SE indicates standard error.

**Table SI3. Average fecundity (oocytes per polyp) for 2014 reproductive dataset.** SE indicates standard error.

**Figure SI1**. Mean percent spermatocyst stage (%) versus treatment including the 2014 reproductive dataset. Sperm stages increase with maturity. N indicates number of individuals and n indicates number of spermatocysts (September 2010: N = 15, n = 1621; June 2011: N = 15, n = 1600; September 2011: N = 8, n = 766; January 2012: N = 14, n = 895; January 2013: N = 15, n = 1149; Time 0: N = 20, n = 2000; Ambient: N = 18, n = 1710; 2100: N = 12, n = 1070).

**Figure SI2. Percent oocyte frequency (%) versus oocyte diameter (μm) by treatment and month.** From top to bottom and left to right: September 2010, June 2011, September 2011, January 2012, January 2013, Time 0, Ambient, and 2100. N indicates number of individuals, n indicates number of oocytes measured, and μ_x_ is the mean, shown by the arrows.

**Figure SI3. Boxplot distribution of fecundity (oocytes per polyp) versus treatment, separated by collection month.** N indicates number of individuals (Time 0: N = 29; June Ambient: N = 10; June 2100: N = 3; September Ambient: N = 20; September 2100: N = 17). **[H2]Figure SI4. Average fecundity (oocytes per polyp) versus treatment, including the 2014 reproductive dataset.** Error bars represent standard error. N indicates number of individuals (September 2010: N = 19; June 2011: N = 19; September 2011: N = 7; January 2012: N = 19; January 2013: N = 13; Time 0: N = 29; Ambient: N = 30; 2100: N = 20; June Ambient: N = 10; June 2100: N = 3; September Ambient: N = 20; September 2100: N = 17).

**SA1. Supplementary appendix 1.** Results of the RMANOVAS and corresponding p-values for the primary text.

**Table SA1_1. OA dataset, Stage I spermatocysts.** H_0_: The percent of stage I spermatocysts does not depend on treatment. ANOVA: p-value = *9.485e-07*

**Table SA1_2. OA dataset, Stage II spermatocysts.** H_0_: The percent of stage II spermatocysts does not depend on treatment. ANOVA: p-value = *6.102e-11*

**Table SA1_3. OA dataset, Stage III spermatocysts** H_0_: The percent of stage III spermatocysts does not depend on treatment. ANOVA: p-value = *2.718e-6*

**Table SA1_4. OA dataset, Stage IV spermatocysts** H_0_: The percent of stage IV spermatocysts does not depend on treatment. ANOVA: p-value = *2.38e-3*

**Table SA1_5. OA dataset, oocyte diameters** H_0_: The oocyte diameter does not depend on treatment. ANOVA: p-value = *<2.2e-16*

**Table SA1_6. OA dataset, fecundity.** H_0_: The fecundity does not depend on treatment. ANOVA: p-value = *7.57e-13*

**SA2. Supplementary appendix 2.** Results of the RMANOVAs and corresponding p-values for the supplementary texts regarding either sprig length, comparison of end month, and comparison to the 2014 reproductive dataset.

**Table SA2_1. All sprig measurement type datasets.** H_0_: The percent change does not depend on measurement type. ANOVA: p-value = *<2.2e-16*

**Table SA2_2. OA dataset with 2014 reproductive dataset, January 2013 spermatocyst stages.** H_0_: The percent of each stage of spermatocyst does not depend on treatment

**Table SA2_3. OA dataset with 2014 Reproductive Dataset, June Ambient to June 2011 Stage II Spermatocysts.** H_0_: The percent of stage II spermatocyst does not depend on treatment

**Table SA2_4. OA dataset with 2014 reproductive dataset, September OA to September 2011 Stage II spermatocysts.** H_0_: The percent of stage II spermatocysts does not depend on treatment. ANOVA: p-value = *1.0e-3*

**Table SA2_5. OA dataset, oocyte diameters: only Time 0 and September 2016.** H_0_: The oocyte diameter does not depend on treatment. ANOVA: p-value = *<2.2e-16*

**Table SA2_6. OA dataset, oocyte diameters: Time 0, Ambient, and 2100 split by month**. H_0_: The oocyte diameter does not depend on treatment. ANOVA: p-value = *<2.2e-16*

**Table SA2_7. OA dataset of oocyte diameters (split by month) with 2014 reproductive dataset.** H_0_: The oocyte diameter does not depend on treatment. ANOVA: p-value = *<2.2e-16*

**Table SA2_8. OA dataset of fecundity by month.** H_0_: The fecundity does not depend on treatment. ANOVA: p-value = *9.357e-5*

**Table SA2_9. OA dataset with 2014 reproductive dataset of fecundity.** H_0_: The fecundity does not depend on treatment. ANOVA: p-value = *1.07e-9*

**Table SA2_10. June OA dataset with June 2011 dataset of fecundity.** H_0_: The fecundity does not depend on treatment. ANOVA: p-value = *4.73e-3*

**Table SA2_11. September OA dataset with September 2010 dataset of fecundity.** H_0_: The fecundity does not depend on treatment. ANOVA: p-value = *1.32e-12*

**SA3. Supplementary appendix 3**. Results of the RMANOVAs and corresponding p-values for the supplementary text regarding the comparison of the sprig datasets with the reproductive data.

**Table SA3_1 OA dataset with sprig length dataset and Stage I spermatocysts**. H_0_: The percent of stage I spermatocysts does not depend on treatment. ANOVA: p-value = 2.27e-1

**Table SA3_2 OA dataset with sprig length dataset and Stage II spermatocysts**. H_0_: The percent of stage II spermatocysts does not depend on treatment. ANOVA: p-value = 0.2948

**Table SA3_3. OA dataset of oocyte diameters with sprig length dataset.** H_0_: The oocyte diameter does not depend on treatment. ANOVA: p-value = *<2.2e-16*

**Table SA3_4. June Ambient dataset of oocyte diameters with June Ambient length dataset.** H_0_: The oocyte diameter does not depend on treatment. ANOVA: p-value = *7.52e-4*

**Table SA3_5. September Ambient dataset of oocyte diameters with September Ambient length dataset.** H_0_: The oocyte diameter does not depend on treatment. ANOVA: p-value = *4.73e-4*

**Table SA3_6. September 2100 dataset of oocyte diameters with September 2100 length dataset.** H_0_: The oocyte diameter does not depend on treatment. ANOVA: p-value = *3.22e-16*

**Table SA3_7. Ambient June dataset of fecundity with Ambient June sprig length dataset.** H_0_: The fecundity does not depend on sprig quality. ANOVA: p-value = 8.9e-1

**Table SA3_8. September Ambient dataset of fecundity with September Ambient sprig length dataset.** H_0_: The fecundity does not depend on sprig quality. ANOVA: p-value = 1

**Table SA3_9. September 2100 dataset of fecundity with September 2100 Sprig length dataset.** H_0_: The fecundity does not depend on sprig quality. ANOVA: p-value = 5.07e-2

